# Modelling the spatial and temporal constrains of the GABAergic influence on neuronal excitability

**DOI:** 10.1101/2021.06.22.449394

**Authors:** Aniello Lombardi, Heiko J. Luhmann, Werner Kilb

## Abstract

GABA (γ-amino butyric acid) is an inhibitory neurotransmitter in the adult brain that can mediate depolarizing responses during development or after neuropathological insults. Under which conditions GABAergic membrane depolarizations are sufficient to impose excitatory effects is hard to predict, as shunting inhibition and GABAergic effects on spatiotemporal filtering of excitatory inputs must be considered. To evaluate at which reversal potential a net excitatory effect was imposed by GABA (E_GABA_^Thr^), we performed a detailed in-silico study using simple neuronal topologies and distinct spatiotemporal relations between GABAergic and glutamatergic inputs.

These simulations revealed for GABAergic synapses located at the soma an E_GABA_^Thr^ close to action potential threshold (E_AP_^Thr^), while with increasing dendritic distance E_GABA_^Thr^ shifted to positive values. The impact of GABA on AMPA-mediated inputs revealed a complex temporal and spatial dependency. E_GABA_^Thr^ depends on the temporal relation between GABA and AMPA inputs, with a striking negative shift in E_GABA_^Thr^ for AMPA inputs appearing after the GABA input. The spatial dependency between GABA and AMPA inputs revealed a complex profile, with E_GABA_^Thr^ being shifted to values negative to E_AP_^Thr^ for AMPA synapses located proximally to the GABA input, while for distally located AMPA synapses the dendritic distance had only a minor effect on E_GABA_^Thr^. For tonic GABAergic conductances E_GABA_^Thr^ was negative to E_AP_^Thr^ over a wide range of g_GABA_^tonic^ values. In summary, these results demonstrate that for several physiologically relevant situations E_GABA_^Thr^ is negative to E_AP_^Thr^, suggesting that depolarizing GABAergic responses can mediate excitatory effects even if E_GABA_ did not reach E_AP_^Thr^.

**Author summary:** The neurotransmitter GABA mediates an inhibitory action in the mature brain, while it was found that GABA provokes depolarizations in the immature brain of after neurological insults. It is, however, not clear to which extend these GABAergic depolarizations con contribute to an excitatory effect. In the present manuscript we approached this question with a computational model of a simplified neurons to determine which amount of a GABAergic depolarizing effect, which we quantified by the so called GABA reversal potential (E_GABA_), was required to turn GABAergic inhibition to excitation. The results of our simulations revealed that if GABA was applied alone a GABAergic excitation was induced when E_GABA_ was around the action potential threshold. When GABA was applied together with additional excitatory inputs, which is the physiological situation in the brain, only for spatially and temporally correlated inputs E_GABA_ was close to the action potential threshold. For situations in which the additional excitatory inputs appear after the GABA input or are distant to the GABA input, an excitatory effect of GABA could be observed already at E_GABA_ substantially negative to the action potential threshold. This results indicate that even slightly depolarizing GABA responses, which may be induced during or after neurological insults, can potentially turn GABAergic inhibition into GABAergic excitation.

## 1. Introduction

The neurotransmitter γ-amino butyric acid (GABA) is the major inhibitory neurotransmitter in the adult mammalian brain [1]. GABA regulates the excitation of neurons and is thus essential for e.g. the control of sensory integration, regulation of motor functions, generation of oscillatory activity, and neuronal plasticity [2–4]. GABA mediates its effects via metabotropic GABA_B_ receptors [5] and ionotropic GABA_A_ receptors, ligand-gated anion-channels with a high Cl^−^ permeability and a partial HCO_3_^−^ permeability [6]. The membrane responses caused by GABA_A_ receptor activation thus depend on the reversal potential of GABA receptors (E_GABA_), which is determined mainly by the intracellular Cl^−^ concentration ([Cl^−^]_i_) and to a lesser extent by the HCO_3_^−^ gradient across the membrane [6].

About 30 years ago seminal studies demonstrated that GABA_A_ receptors can mediate depolarizing and excitatory actions in the immature central nervous system [7–9]. This depolarizing GABAergic action reflects differences in the [Cl^−^]_i_ homeostasis between immature and adult neurons [10–15]. In particular, low functional expression of a K^+^-Cl^−^ cotransporter (KCC2), which mediates the effective extrusion of Cl^−^ and thus establishes the low [Cl^−^]_i_ required for hyperpolarizing GABAergic membrane responses [16], prevent hyperpolarizing GABA responses in the immature brain. In addition, the inwardly directed Cl^−^ transporter NKCC1 mediates the accumulation of Cl^−^ above passive distribution that underlies the depolarizing membrane responses upon activation of GABA_A_ receptors [17–21]. These depolarizing GABAergic membrane responses play a role in several developmental processes [11,22,23], like neuronal proliferation [24], apoptosis [25], neuronal migration [26], dendro- and synaptogenesis [27], timing of critical periods [28] and the establishment of neuronal circuitry [29]. Of clinical importance, an elevated [Cl^−^]_i_ is also a typical consequence of several neurological disorders in the adult brain, like trauma, stroke or epilepsy and is considered to augment the consequences of such insults [11,30,31].

However, it is important to consider that depolarizing GABA responses do not per se lead to excitatory effects. In fact, the membrane shunting that unescapably accompanies the activation of GABA_A_ receptors can dominate over the excitatory effects of the membrane depolarization [11,32–34]. Theoretical considerations [35,36] suggest that the relation between E_GABA_ and the action potential threshold (E_AP_^Thr^) determine whether activation of GABA_A_ receptors mediates excitatory (E_GABA_ positive to E_Thr_^AP^) or inhibitory (E_GABA_ negative to E_Thr_^AP^) actions. However, this concept is probably an oversimplification, as within the dendritic compartment the local activation of GABAergic conductance influences not only the amplitude of local excitatory synaptic postsynaptic potentials (EPSPs), but also the length and time constants of the dendritic membrane and thus temporal and spatial summation of excitatory synaptic inputs [37,38]. Moreover, the depolarizing effect of GABAergic stimulation outlasts the conductance increase associated with GABA_A_ receptor activation, resulting in a bimodal GABA effect. Close to the initiation of GABAergic responses the shunting effect of the enhanced GABAergic conductance dominate and mediate inhibition. This phase is followed by an excitatory phase dominated by the GABAergic depolarization [39,40]. In addition, E_Thr_^AP^ is a dynamic variable, that depends on the background conductance and the density and adaptation state of voltage-gated Na^+^ channels [10,41,42]. Experimental studies on the effects of GABAergic inputs on neuronal excitability demonstrated for immature neocortical neurons that E_GABA_ required for excitatory GABAergic responses (E_GABA_^Thr^) was close to E_AP_^Thr^ [43], while in immature hippocampal neurons E_GABA_^Thr^ was considerably negative to E_AP_^Thr^ [44]. The observations that (i) the GABA effect can switch from inhibition to excitation for delayed glutamatergic inputs [39], that (ii) GABA inputs in distal dendrites can facilitate neuronal excitability [40], and that (iii) extrasynaptic GABAergic activation mediates an excitatory effect whereas synaptic inputs mediate inhibition [45], also suggest that the reversal potential required for GABAergic excitation is not only defined by E_AP_^Thr^. This complexity is further supported by recent in-vivo investigations that identified excitatory as well as inhibitory effects of GABA in the immature brain [46–48]. In summary, to our knowledge no clear concept is currently available that can explain how E_GABA_ influences GABAergic excitation/inhibition and the effect of GABA on spatiotemporal summation of EPSPs in the dendritic compartment.

Therefore, the present computational study investigates the dependency between E_GABA_ and excitatory and inhibitory consequences of GABA_A_ receptor activation and attempts to establish a general view of the impact of depolarizing GABAergic effects on the excitability of neurons. Our results demonstrate that only for GABAergic synapses located at or close to the soma the difference between E_GABA_ and E_AP_^Thr^ predicts whether GABA has an excitatory or an inhibitory action. The E_GABA_ at which depolarizing GABA actions switch from inhibition to excitation is in most cases negative to E_AP_^Thr^ and depends on the temporal and spatial relation between GABA and AMPA inputs, with a more excitatory effect on AMPA inputs that are delayed or located proximal to GABA inputs. We conclude from our results that GABA can mediate excitatory effects even if E_GABA_ is considerably hyperpolarized to E_AP_^Thr^.

## 2. Results

### 2.1. Simulation of active and passive properties of immature CA3 pyramidal neurons

The parameters used for the models in this study are based on the cellular properties obtained in whole-cell patch-clamp recordings from visually identified CA3 neurons in horizontal hippocampal slices from P4-7 mice. Some parameters of these recordings have been used in our previous report [49]. The analysis of the patch-clamp experiments revealed that the immature CA3 pyramidal neurons had an average resting membrane potential (RMP) of −50.5 ± 1.3 mV, an average input resistance (R_Inp_) of 1.03 ± 0.11 GOhm, and an average membrane capacity (C_M_) of 132.3 ± 33.6 nF (all n=42). As the passive membrane properties directly influence synaptic integration as well as active properties, like E_AP_^Thr^ or the shape of the action potential (AP), we first adapted the spatial properties and the passive conductance g_pas_ of the ball-and-stick model to emulate the recorded properties. To obtain sufficient similarity for these parameters between the model and the real cells we equipped a ball-and-stick model (soma diameter (d) = 46.6 μm, dendrite length = 1 mm, dendrite diameter = 1 μm) with a passive conductance density (g_pas_) of 1.28*10^−5^ nS/cm^2^ at a reversal potential (E_pas_) of −50.5 mV. This model had a RMP of −50.5 mV, a R_Inp_ of 1.045 GOhm and a C_M_ of 144.4 nF. In some experiments we reduced the topology to a simple ball model (*one node, d =* 46.6 μm), without adapting g_pas_, to evaluate the impact of GABA under quasi one-dimensional conditions.

With these configurations we next implemented a mechanism that provided APs with properties comparable to the APs recorded in CA3 pyramidal neuron. In particular, we were interested to simulate the AP properties around AP initiation as precisely as possible, because for the main questions of this manuscript we are interested in the E_AP_^Thr^. Since it was not possible to generate a reasonable sharp AP onset with a standard Hodgkin-Huxley (HH) model, we used a modified Markov model (see materials for details) to simulate AP with a considerable precision (Fig. 1A-E).

**Figure 1.**
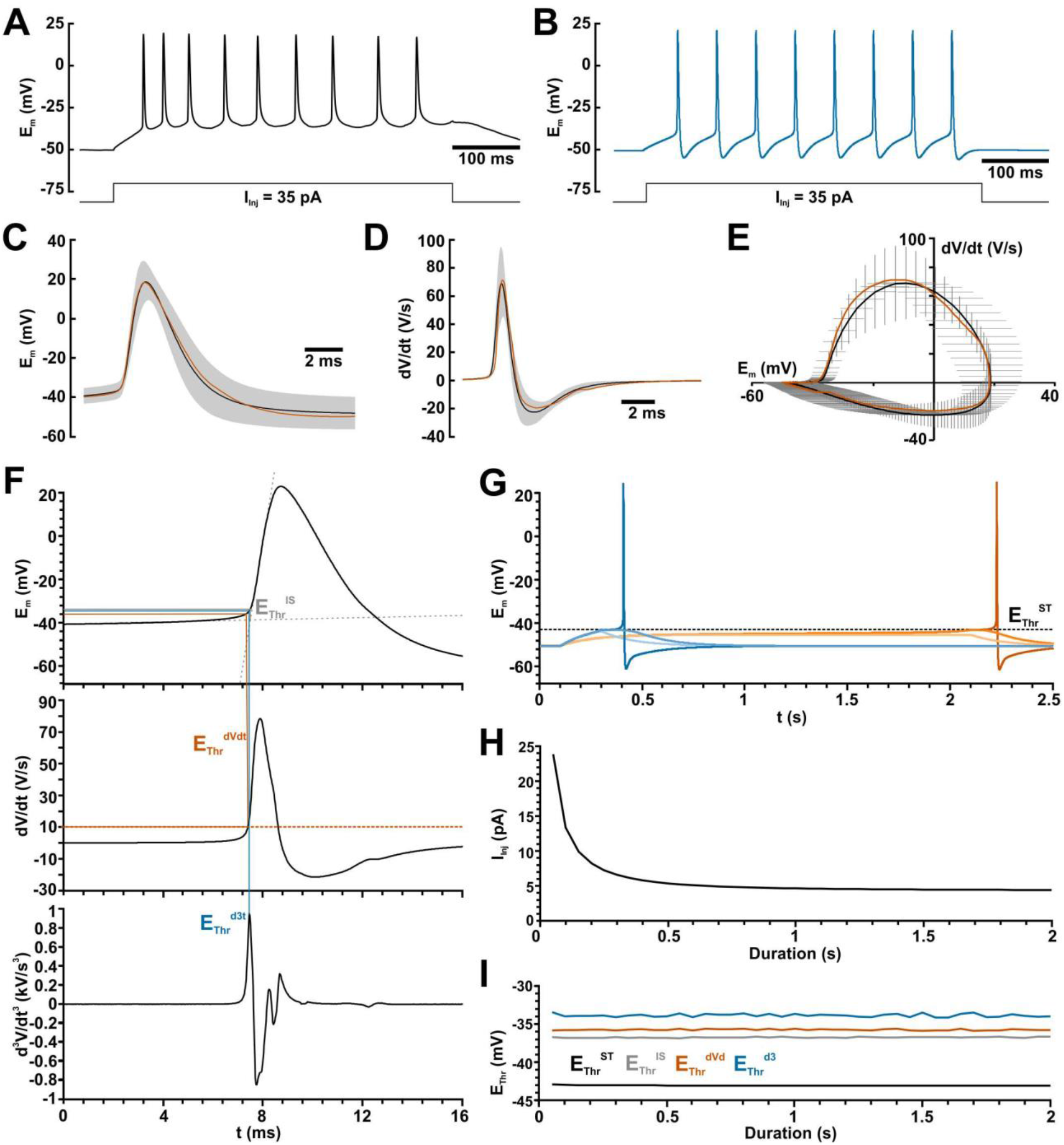
Properties of recorded and simulated action potentials (APs). A: Typical AP train recorded in a CA3 pyramidal neuron upon a current injection of +35 pA. B: AP train simulated in a ball-and-stick model using the modified Markov model. C: Average voltage trace of recorded APs (black line = average; gray area ± SEM) and of the simulated AP (orange trace). D: Discharge rate of recorded (black line, gray area) and simulated AP (orange trace). E: Phase plane plot of recorded (whiskers = mean ± SEM) and simulated AP (orange trace). F: Determination of the AP threshold from the intersection of linear voltage fits (E_Thr_^IS^, gray lines), from the time point dV/dt reaches the 10 V/s threshold (E^Thr^_dVdt_, orange lines), and from the time point d^3^V/dt^3^ reaches the peak value (E_Thr_^d3^, blue lines). G: Determination of the AP threshold at maximal potential of a subthreshold depolarization (E_Thr_^ST^, black lines). Blue traces indicate a 200 ms depolarizing stimulus and orange traces a 2 s stimulus. Dark tones indicate the smallest suprathreshold stimulus, middle tones the largest subthreshold stimulus and light tones a clearly subthreshold stimulus. H: Injection current (I_Inj_) required to elicit an AP at different stimulus durations. I: Values of different AP threshold parameters for various stimulation durations. Note that AP threshold is independent from the stimulation duration.

Because the relation between E_AP_^Thr^ and E_GABA_ is one major parameter investigated in this study and since no clear definition of the AP threshold has been given [42], we initially used 4 different methods to determine the action potential threshold (Fig. 1F): 1.) The AP threshold value E_Thr_^dVdt^ was defined as the potential at which dV/dt first crosses a velocity of 10 V/s [44,50] (Fig. 1F orange lines). 2.) E_Thr_^d3^ was defined as the potential at the time point of the first positive peak in d^3^V/dt^3^ [51] (Fig. 1F, blue lines). 3.) E_Thr_^IS^ was determined at the intersection between linear regressions of the baseline before the AP and the rising phase of an AP (Fig. 1H) [43] (Fig. 1F, gray lines). 4.) E_Thr_^ST^ was defined as the maximal potential reached at the strongest subthreshold stimulation (Fig. 1G, dashed line), i.e. the minimal potential that did not lead into the regenerative Hodgkin cycle. While the rheobase, i.e. the minimal suprathreshold injection current, demonstrated as expected a hyperbolic increase at shorter stimulus durations and converged to 4.4495 pA (Fig. 1H), the distinct E_AP_^Thr^ parameters are virtually independent on the duration of the stimulus (Fig. 1I). In the ball model average E_Thr_^dVdt^ was −35.7 mV, average E_Thr_^d3^ was −33.8 mV, average E_Thr_^IS^ was −36.7 mV, and E_Thr_^ST^ converged to −43.04 mV (Fig. 1I). When using the ball-and-stick model the rheobase was slightly larger at 6.708 pA, E_Thr_^dVdt^ was −35.7 mV, E_Thr_^d3^ was −33.8 mV, E_Thr_^IS^ was −36.6 mV, and E_Thr_^ST^ converged to −42.71 mV (data not shown).

Because for the following simulations several hundred sweeps were required for each analyzed parameter and thus a time-effective simulation was compulsory, we next evaluated the maximal dt interval required to obtain stable AP responses. This experiment demonstrated that the time course of AP and E_AP_^Thr^ determination remained stable until a dt of 0.1 ms (Suppl. Fig. 1). Thus we decided to use a dt of 0.1 ms in the following simulations.

### 2.2. Determination of the threshold for excitatory GABAergic responses

To identify the reversal potential at which the GABA response switches from inhibitory to excitatory, we first determined the GABAergic conductance that was sufficient to trigger an AP, which was defined as the GABAergic excitation threshold (g_GABA_^Thr^). The value of g_GABA_^Thr^ was determined by systematically increasing the conductance of a simulated GABAergic input until an AP was evoked. To determine this excitation threshold as precisely as possible, we used a multi-step procedure to incrementally confine the threshold conductance (Fig. 2A). This procedure was repeated for a whole set of E_GABA_ values (Fig. 2B).

**Figure 2.**
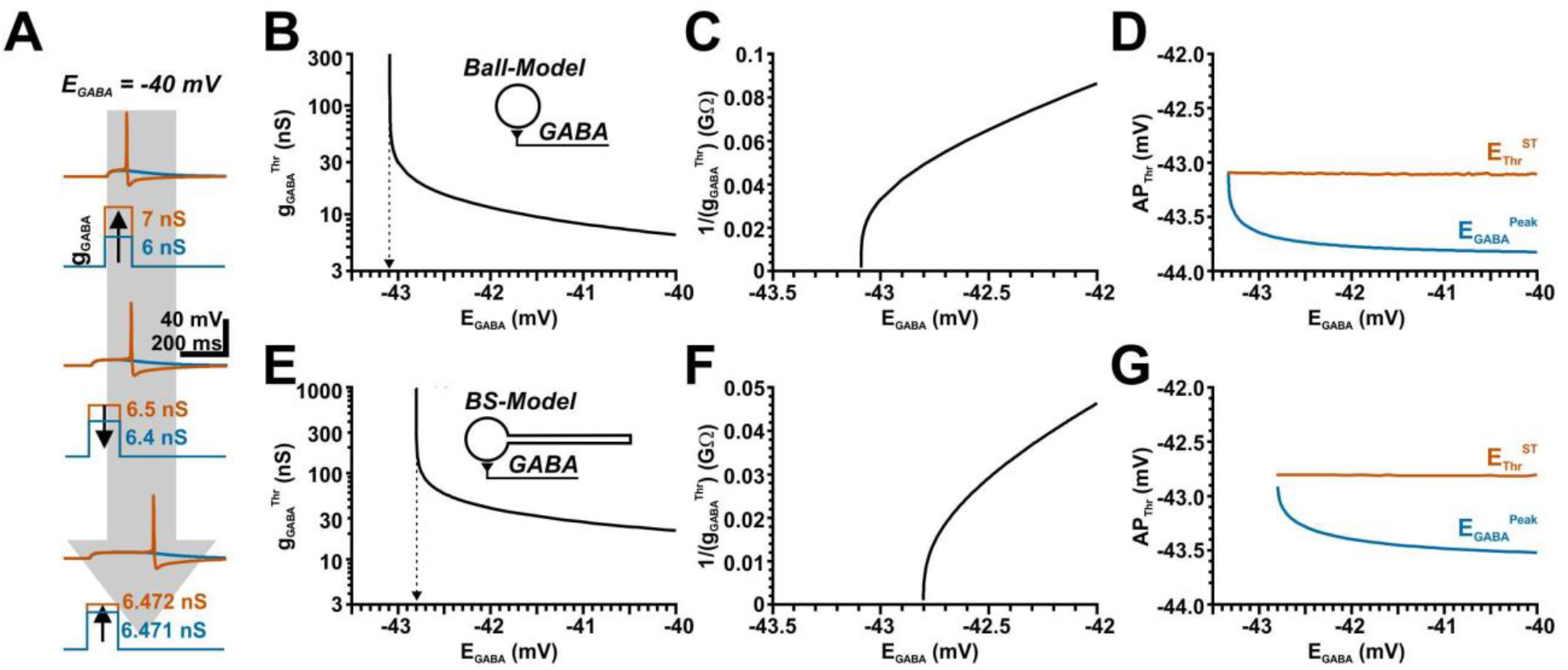
Determination of the threshold conductance at different E_GABA_ enable the identification of E_GABA_ at which responses switch from inhibitory to excitatory (E_GABA_^Thr^). A: Typical voltage traces illustrating the mechanisms used to determine the threshold g_GABA_ value. For this purpose, gGABA was increased until the first AP was induced (upper panel), then decreased by finer g_GABA_ steps until the AP disappears (middle panel), followed by a subsequent increase in g_GABA_ with finer g_GABA_ steps (lower panel). In total, 6 alternating rounds of increased/decreased g_GABA_ steps were used. The g_GABA_ value required to induce an AP in the last increasing step was considered as threshold (g_GABA_^Thr^). B: Plotting g_GABA_^Thr^ versus E_GABA_ demonstrate that with decreasing E_GABA_ higher g_GABA_^Thr^ values were required, which approximated infinite values. C: A reciprocal plot of g_GABA_^Thr^ enables the precise determination of E_GABA_^Thr^. At E_GABA_ values negative to E_GABA_^Thr^ no action potential could be induced, suggesting a stable GABAergic inhibition. D: The determined AP threshold E_Thr_^ST^ (orange line) is constant over various E_GABA_, whereas the peak potential of the GABAergic depolarization, which was determined at g_GABA_^Thr^ in absence of AP mechanism (E_GABA_^Peak^, blue line) increases with decreasing E_GABA_. Note that the values converged in one point when E_GABA_ reaches E_Thr_^ST^. E-G: Similar plots for a ball-and-stick model. Note that E_GABA_^Thr^ was shifted to less negative values in this configuration.

In the ball model (*one node, d =* 46.6 μm) these systematic simulations demonstrated an obvious hyperbolic increase of g_GABA_^Thr^ when E_GABA_ approaches values below −43 mV (Fig. 2B). The g_GABA_^Thr^ curve approximated an E of −43.09 mV, which was precisely determined from a reciprocal plot of the g_GABA_^Thr^ values (Fig. 2C). Negative to an E_GABA_ of −43.09 mV no action potential could be evoked, regardless of the amount of g_GABA_. These E_GABA_ values thus reflects the threshold, at which GABA actions can mediate a direct excitation and we termed this value “threshold E_GABA_” (E_GABA_^Thr^). Note that this value is close to the E_Thr_^ST^ value of −43.04 mV determined in the previous experiments. Since E_AP_^Thr^ is influenced directly by the total membrane conductance, we also determined the amplitude of the GABAergic voltage response under conditions when the AP initiation was blocked (E_GABA_^Peak^) as well as different E_AP_^Thr^ parameters. These analyses revealed that E_Thr_^d3^ was around −29 mV for all E_GABA_. E_Thr_^ST^ was relatively stable around −43.1 mV, with a slight positive shift at low E_GABA_ values that converges to −43.09 mV (Fig. 2D). E_GABA_^Peak^ was for higher E_GABA_ around −43.8 mV and showed a positive shift with decreasing E_GABA_ that converged to values of −43.1 mV (Fig. 2D).

In summary, these results indicate that GABA acts as excitatory neurotransmitter as long as E_GABA_ is positive to −43.09 mV, which is extremely close to the AP threshold E_Thr_^ST^. This observation is in line with previous predictions that propose exactly this relation between E_AP_^Thr^ and E_GABA_ [35,36]. In addition, our simulations suggest that E_Thr_^ST^ is probably the most relevant definition for E_AP_^Thr^ if the direction of GABA effects should be predicted from the difference between E_GABA_ and E_AP_^Thr^.

Next we performed the same simulation with a ball-and-stick model. These simulations revealed that the g_GABA_^Thr^ curve approximated an E_GABA_ of −42.8 mV (Fig. 2E-F), which is in the range of the E_Thr_^ST^ (−42.71 mV) determined for the ball-and-stick model. E_Thr_^d3^ was around −29.8 mV for all E_GABA_. E_Thr_^ST^ was stable at values around −42.8 mV and converges at low E_GABA_ to −42.8 mV (Fig. 2G). E_GABA_^Peak^ was for higher E_GABA_ around −43.6 mV and converged with decreasing E_GABA_ to −42.8 mV (Fig. 2G). Thus, E_GABA_^Thr^ for a somatic synapse is still in good agreement with the AP threshold value E_Thr_^ST^ with this slightly more complex neuronal topology.

For the next set of experiments, we located a single GABA synapse along the dendrite of the ball- and-stick model and determined E_GABA_^Thr^ for each of these 20 synaptic positions, using the method described above. The considerable conductance and capacitance provided by the dendritic membrane leads, as expected, to a reduced amplitude and a slower time course of the GABAergic PSPs recorded at the dendritic positions (Fig. 3A). Accordingly, larger g_GABA_ values were required to trigger APs for more distant dendritic locations of GABAergic inputs (Fig. 3B, C). At the most distant dendritic positions gGABA values above 100 nS (i.e. more than 100x of g_GABA_ of a single synaptic event [49]) were required to trigger an AP, which virtually clamped the dendritic membrane at the synapse position to E_GABA_ (Fig. 3B). A systematic analysis of g_GABA_^Thr^ at different E_GABA_ values illustrated that g_GABA_^Thr^ showed a considerable less steep dependency on E_GABA_ at more distant dendrite positions (Fig. 3C). The reciprocal plot of g_GABA_^Thr^ demonstrated that the g_GABA_^Thr^ values did not converge at similar E_GABA_ values for the different synapse locations, but that the curves reached the abscissa at considerable more positive values for distant GABAergic inputs (Fig. 3D). Intriguingly, E_GABA_^Thr^ values were close to E_Thr_^ST^ for synapses close to the soma, were shifted to slightly more negative E_GABA_^Thr^ values for dendritic synapses at a distance of ca. 250 μm, and increased to more positive E_GABA_^Thr^ values with additional distance to the soma (Fig. 3E). E_GABA_^Peak^, which was determined in the absence of AP mechanisms and reflects the effective voltage fluctuation at the soma and thus the AP initiation site, was shifted to negative potentials at more distant dendritic positions (Fig. 3G, H), while the position of GABA synapses had no major effect on E_Thr_^ST^ (Fig. 3F, H). In summary, these simulations revealed that E_GABA_^Th^ is not close to the AP threshold value E_Thr_^ST^ for synapses that are located in the dendrite, but that E_GABA_^Th^ is shifted to more positive values with increasing distance. This observation suggests that for dendritic synapses a more positive E_GABA_ (corresponding to a higher [Cl^−^]_i_) is required to mediate a direct excitatory effect.

**Figure 3.**
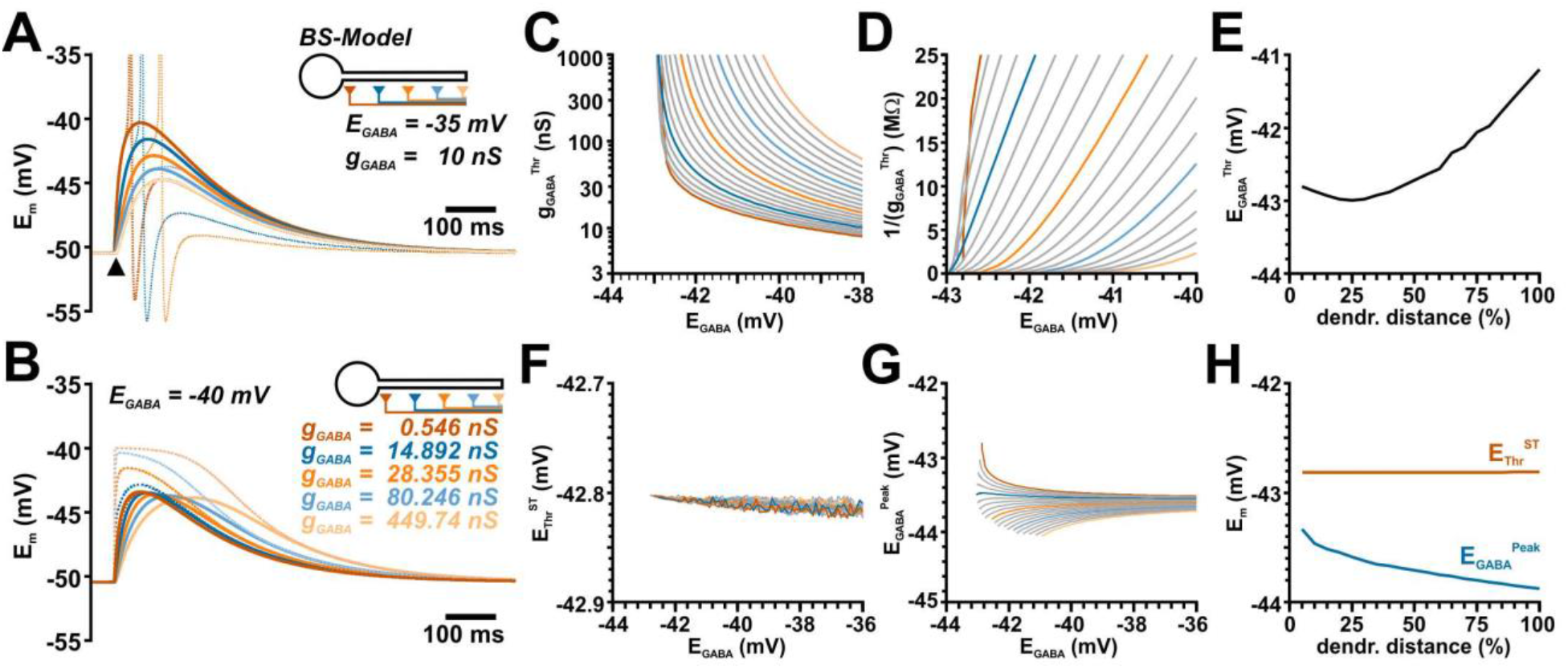
Determination of E_GABA_^Thr^ at different dendrite positions. A: Simulated voltage traces obtained with the given parameters at different locations as indicated by color code. The dashed traces represent simulation with added AP mechanism. The amplitude of GABA responses clearly depends on the dendritic location. B: Simulated voltage traces for g_GABA_^Thr^ and E_GABA_ of −40 mV at the soma (solid trace) and the synaptic site (dashed line). Color codes the different synapse locations. For each location different g_GABA_ (as indicated) had to be used. Note that at distant synapses considerable large g_GABA_ were required, which virtually clamped E_m_ at the synaptic site to E_GABA_. C: Systematic plot of g_GABA_^Thr^ determined at various E_GABA_. The curves were obtained from 20 equidistant positions along the dendrite. The 1^th^, 5^th^, 10^th^, 15^th^ and 20^th^ trace is color-coded as in A for better readability. D: The reciprocal plot of g_GABA_^Thr^ revealed that the curves did not monotonically approach the abscissa. Therefore, E_GABA_^Thr^ was estimated from a linear fit to the last two data-points. E: E_GABA_^Thr^ showed a considerable shift towards depolarized potentials with increasing dendritic distance. F: The AP threshold E_Thr_^ST^ remained rather stable with different E_GABA_ or different synaptic location. G: The peak potential (E_GABA_^Peak^) of the somatic GABAergic depolarization at g_GABA_^Thr^ converges toward E_Thr_^ST^ only for soma-near synapses (dark orange trace). With more distant synapses less depolarized E_GABA_^Peak^ was required. Color code as in C. H: While the average E_Thr_^ST^ (orange line) is stable for all dendritic locations, the average E_GABA_^Peak^ (blue line) is shifted to more negative values with increasing dendritic distance.

### 2.3. Effect of phasic GABAergic inputs on glutamatergic excitation

The previous results demonstrated that only at perisomatic synapses E_GABA_^Thr^ was reached when E_GABA_ was at the action potential threshold E_Thr_^ST^, but that E_GABA_^Thr^ was systematically shifted to positive E_GABA_ at distant synapses in a ball-and-stick model. However, these experiments do not reflect the physiological situation of GABAergic transmission in the brain. First, the threshold conductance g_GABA_^Thr^ determined by these simulations is above physiological values for moderate GABAergic inputs [49,52,53] making a direct excitatory GABAergic input implausible. And second, synaptic activity is characterized by the co-activation of GABA and glutamate receptors [54–56], with the latter constituting the main excitatory drive [57]. Therefore, we next simulated the impact of a GABAergic co-stimulation on glutamatergic synaptic inputs and determined the g_AMPA_ values that were required to trigger an AP. As in the previous experiments, we varied E_GABA_ to determine E_GABA_^Thr^, which is defined as the E_GABA_ value at which the GABAergic effect shifts from inhibitory (i.e. GABA co-activation requires larger g_AMPA_ to trigger APs) to excitatory action (i.e. GABA co-activation requires less g_AMPA_) (Fig. 4A). This effect was quantified as the GABAergic excitability shift (Δg_AMPA_^Thr^), with g_AMPA_^Thr^ describing the g_AMPA_ value sufficient to trigger an AP, and Δg_AMPA_^Thr^ defined as difference in g_AMPA_^Thr^ between conditions with and without GABAergic co-stimulation [Δg_AMPA_^Thr^ = (g_AMPA_^Th^)_withGABA_ − (g_AMPA_^Th^)_w/oGABA_].

**Figure 4.**
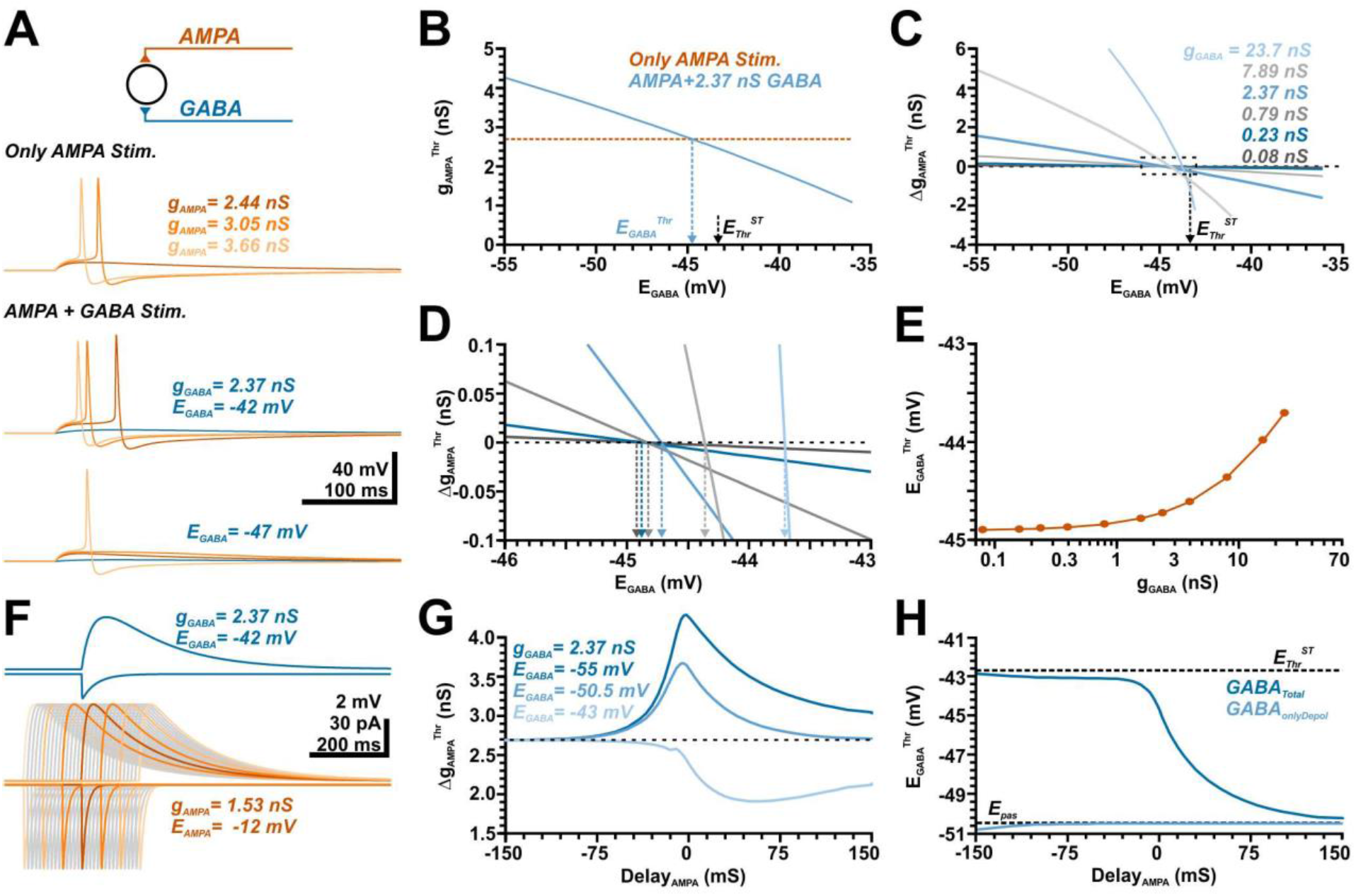
Influence of a GABAergic input at different E_GABA_^Thr^ on the AMPA receptor-dependent excitation threshold. A: Simulated voltage traces illustrating the membrane responses induced by three different conductances of the AMPA synapse in the absence (top traces) and the presence of a simultaneous GABAergic input at E_GABA_ of −42 mV (middle traces) and −47 mV (lower traces). B: Plot of the minimal gAMPA required to trigger an AP (g_AMPA_^Thr^) versus the E_GABA_ of the synchronous GABA input (g_GABA_ = 2.37 nS). The E_GABA_ value at which this curve intersects with g_AMPA_^Thr^ determined in the absence of GABA (orange line) defines the GABA concentration at which GABA switches from excitatory to inhibitory (E_GABA_^Thr^). C: Plot of Δg_AMPA_^Thr^ versus E_GABA_ for different g_GABA_ values, as indicated in the graph. D: A magnification of the marked area in C allows the determination of E_GABA_^Thr^ for the different g_GABA_, color code as indicated in C. E: Plot of the E_GABA_^Thr^ determined at different g_GABA_. Note that E_GABA_^Thr^ is substantially negative to E_Thr_^ST^ and increases at higher g_GABA_. F: Simulation of membrane currents (downward deflections) and membrane changes (upward deflections) upon a GABAergic (blue traces) and glutamatergic stimulation. Gray and light orange traces represent temporally shifted glutamatergic inputs, performed to differentiate the effects of conductance vs. depolarization effects. Note that the depolarization shift outlasts the conductance shift for both inputs. G: Influence of the timing between AMPA and GABA input on Δg_AMPA_^Thr^ determined at 3 exemplary E_GABA_. Note that the maximal inhibitory effect at hyperpolarizing (E_GABA_ = −55 mV) or pure shunting GABAergic inputs (E_GABA_ = −50.5 mV) were observed for synchronous AMPA inputs, while the excitatory influence of GABA at depolarized E_GABA_ of −43 mV was maximal for substantially delayed AMPA inputs. H: Quantification of E_GABA_^Thr^ (dark blue) for different delays between GABA and AMPA inputs. Note that for AMPA inputs preceding GABA inputs E_GABA_^Thr^ was close to the AP threshold, while for AMPA inputs lagging GABA inputs E_GABA_^Thr^ approximated −50.5 mV. The light blue traces represent E_GABA_^Thr^ determined for pure simulated GABAergic depolarizations which persistently results in a E_GABA_^Thr^ close to −50.5 mV.

In the first set of experiments we simulated the effect of GABA pulses provided synchronously with AMPA inputs in a ball model (Fig. 4A) using a constant g_GABA_ of 2.37 nS. These experiments demonstrated that the co-stimulation of a GABAergic input can attenuate or enhance AP triggering upon glutamatergic stimulation, depending on E_GABA_ (Fig. 4A). As expected, such a GABA co-stimulation enhanced g_AMPA_^Thr^ at hyperpolarized E_GABA_, while smaller g_AMPA_^Thr^ values were required at more depolarized E_GABA_ (Fig. 4B). From the intersection of this g_AMPA_^Thr^ with the g_AMPA_^Thr^ recorded in the absence of GABAergic inputs we determined that E_GABA_^Thr^ amounted to −44.7 mV under this condition (Fig. 4B), which is considerable more negative than E_Thr_^ST^ of −43.04 mV in the ball model. Additional experiments with different g_GABA_ values revealed that E_GABA_^Thr^ depends on g_GABA_ (Fig. 4C-E). However, only at rather large g_GABA_ values E_GABA_^Thr^ approached toward values > −44 mV. At lower, physiologically more relevant g_GABA_ values E_GABA_^Thr^ converges to a value of −44.9 mV (Fig. 4E). This observation indicates that E_GABA_^Thr^ was consistently lower than E_Thr_^ST^, implying that GABAergic inputs are under these conditions more excitatory than expected from the difference between E_GABA_ and E_AP_^Thr^.

Is has already been proposed that the GABAergic depolarization outlasts the GABAergic currents and can add an additional excitatory drive to neurons [39]. Our simulations replicated this typical behavior, both GABAergic and glutamatergic membrane depolarization outlasted the time course of the respective currents (Fig. 4F). To investigate whether the systematic shift of E_GABA_^Thr^ towards more hyperpolarized potentials was indeed caused by the differential impact of GABAergic conductance and GABAergic membrane depolarization on the AMPA-mediated excitation, we systematically advanced or delayed the time point of AMPA inputs (Fig. 4 F). These simulations revealed that, as expected, the strongest inhibitory effect of a GABAergic input for both hyperpolarizing (at E_GABA_ < RMP) and shunting inhibition (at E_GABA_ = RMP) was observed when it was synchronous to the glutamatergic input (Fig. 4G). In contrast, at more depolarized E_GABA_ the maximal excitatory effect occurred when the AMPA input was given about 60 ms after the GABA input (Fig. 4G, light trace), i.e. at a time point when the GABAergic conductance virtually ceased, but a considerable GABAergic depolarization persisted (Fig. 4F, blue traces). A systematic determination of E_GABA_^Thr^ for different delays demonstrated that E_GABA_^Thr^ was relatively stable around −43 mV for APMA inputs that preceded GABA inputs, and was thus close to E_Thr_^ST^ (Fig. 4H). In contrast, with increasing delays of the glutamatergic inputs E_GABA_^Thr^ converged to −50.5 mV, i.e. to the RMP determined by the reversal potential of the passive membrane conductance (Fig. 4H). In summary, these findings suggest (i) that at preceding AMPA inputs the influence of GABA on this glutamatergic input was dominated by the GABAergic conductance change and thus converged to E_Thr_^ST^ and (ii) that at delayed glutamatergic inputs the influence of GABA on this glutamatergic input was dominated by the GABAergic depolarization.

In the absence of a GABAergic conductance shift each depolarization above −50.5 mV should reduce the distance to the E_AP_^Thr^ and should thus impose an excitatory effect. To verify this hypothesis, we recorded the GABAergic currents at different E_GABA_ and replayed these currents to the modelled neurons via I-clamp, thereby isolating the effect of the GABAergic depolarization from the conductance shift. Indeed, these simulations demonstrated that the effect of the pure GABAergic depolarization reversed at an E_GABA_ of −50.5 mV (Fig. 4H, light trace).

In summary these experiments demonstrated that the effect of a GABAergic stimulus on glutamatergic synaptic inputs cannot simply be predicted from the difference between E_GABA_ and the E_AP_^Thr^ threshold, but that, depending on the temporal relation between GABAergic and glutamatergic inputs, E_GABA_ is substantially lower than E_AP_^Thr^ and thus GABA acts more excitatory than expected from the E_GABA_ to E_AP_^Thr^ relation.

In the next set of experiments, we evaluated how the spatial relation between GABAergic and glutamatergic inputs affects E_GABA_^Thr^ in a ball-and-stick model. For these simulations, we systematically varied both, GABA and AMPA synapse along the dendrite, using 20 equidistant positions each (Fig. 5A), and stimulated both synapses.

**Figure 5.**
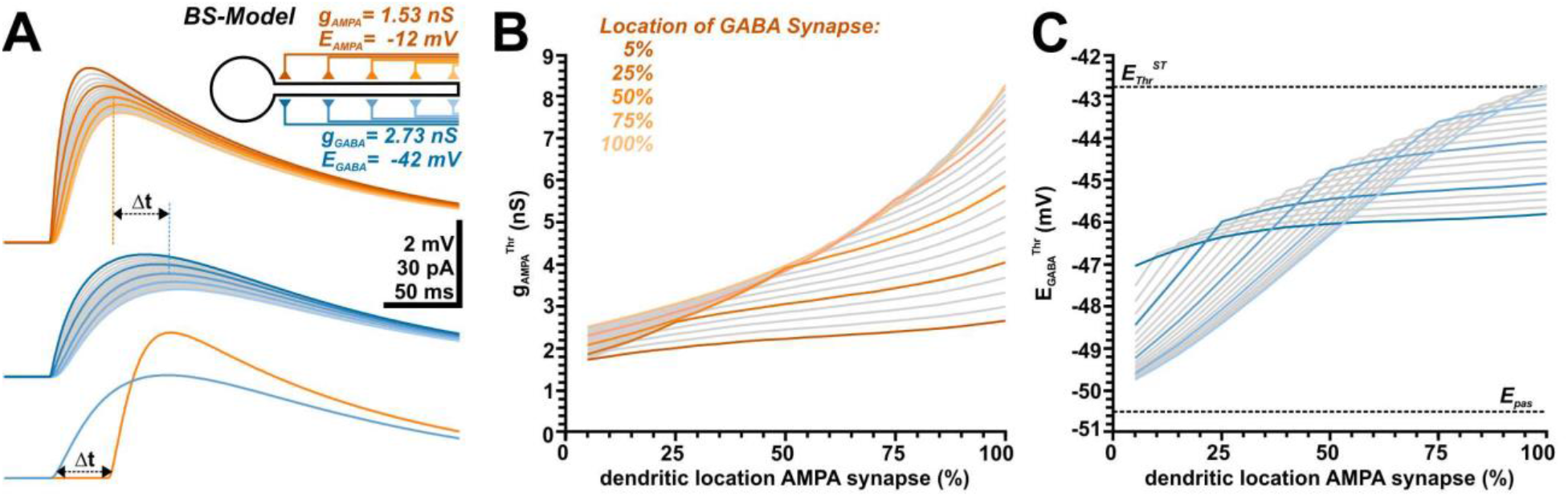
Influence of the spatial relation between the AMPA receptor-dependent and the GABA receptor-dependent synaptic input on g_AMPA_^Thr^ and E_GABA_^Thr^. A: Simulated voltage traces illustrating the membrane responses induced by AMPA synapses (orange traces) and by GABA synapses (blue traces) located at different dendritic locations. The colored traces represent synapses at 5%, 25%, 50%, 75% and 100% of the dendritic length, as color-coded in the schematic inset. Note the slower onset kinetics and delayed peak for distant dendritic synapses. The lower traces depict how the delay of GABA and AMPA was adjusted to obtain synchronous peak depolarizations. B: Effect of the dendritic location on g_AMPA_^Thr^ simulated for 20 equidistant positions of the GABAergic synapse (g_GABA_ =7.89 nS; E_GABA_ = −44 mV). Each line represents the results for one GABA synapse position, the color code identifies every 5^th^ position as indicated. Note the shallow dependency of Δg_AMPA_^Thr^ for proximal and the steep dependency for distal GABA synapses. C: Dependency of E_GABA_^Thr^ on the dendritic positions of AMPA synapses, each line represents the results for one GABA synapse position, with shade coding as in B. Note the shallow location dependency with E_GABA_^Thr^ around −46 mV for the proximal GABA synapses, while for distal GABA synapses a steep E_GABA_^Thr^ profile between ca. −43 mV and −50 mV was observed.

Simulations of single inputs revealed that the time course of the glutamatergic and GABAergic depolarizations critically depended on the dendritic location (Fig. 5A), which reflect spatial filtering [58]. To prevent that this temporal scattering affects the spatial analysis of GABA/AMPA relations, we determined the maximum of the depolarization in control sweeps performed before each run of the definite simulation for each combination of g_AMPA_, AMPA location, E_GABA_, and GABA location in the absence of an AP mechanism. For the definite simulation sweep the temporal relation between glutamatergic and GABAergic input was shifted such that peak depolarization of GABA and AMPA responses coincided (Fig. 5A).

To get an impression how a depolarizing GABAergic input at different locations influences g_AMPA_^Thr^, we first varied the position of a GABAergic synapse with a g_GABA_ of 7.89 nS and an E_GABA_ of −44 mV along the dendrite and determined g_AMPA_^Thr^ for each of the 20 AMPA synapse along the dendrite (Fig. 5B). These simulations showed, as expected, that (i) g_AMPA_^Thr^ increased with increasing dendritic distance, and (ii) that for a soma-near GABAergic synapse the excitatory effect of GABA was stronger than for distal dendritic locations, as indicated by the larger g_AMPA_^Thr^ required for the distal GABA synapses (Fig. 5B). However, we could also demonstrate that (iii) the slope of the g_AMPA_^Thr^ became shallower for AMPA inputs distal to the GABA inputs (Fig. 5B), indicating a strong non-linear influence of GABAergic inputs. To determine how the spatial relation between glutamatergic and GABAergic inputs affects E_GABA_^Thr^ we subsequently varied E_GABA_ (at g_GABA_ of 7.89 nS) for all combinations of synaptic positions and determined when Δg_AMPA_^Thr^ switches the direction (Fig. 5C). These simulations revealed a complex relation between these three parameters. If the GABAergic synapse was located in the proximal dendrite close to the soma (Fig. 5, dark trace) E_GABA_^Thr^ was only weakly dependent on the site of the AMPA synapse and amounted to values between ca. −45 mV and −46 mV. If the GABA synapse was located more distally (Fig. 5, lighter trace) E_GABA_^Thr^ showed a step dependency on the location of the AMPA synapse for all AMPA synapses located proximally to the GABA synapse, while the shallow dependency was maintained for the more distal synapses (Fig. 5C). Under this condition E_GABA_^Thr^ approached −50 mV for proximal AMPA synapses, i.e. when both synapses were 950 μm apart and thus the GABAergic depolarization dominated over the more local shunting effect (see lightest blue trace in Fig. 5C). In contrast, for most distally located AMPA and GABA synapses, which represent spatially correlated inputs distant from the AP initiation zone, E_GABA_^Thr^ approached E_Thr_^ST^ (Fig. 5C).

In summary, these results demonstrate that both, the spatial relation between GABAergic and glutamatergic synapses as well as the location of the GABA synapse influences E_GABA_^Thr^. However, only for spatially correlated inputs at distal dendrites E_GABA_^Thr^ was close to the E_AP_^Thr^. With increasing distance between both synapses and with a closer approximation of the GABA synapse to the soma, E_GABA_^Thr^ was shifted to more negative values, again indicating that GABA mediates a more prominent excitatory action than expected from the difference between E_GABA_ and E_AP_^Thr^.

### 2.4. Effect of tonic GABAergic inputs on glutamatergic excitation

GABA influences neuronal excitability not only via synaptic inputs, but also extrasynaptic, tonic GABAergic currents substantially contribute to the GABAergic effects [59,60] and can mediate even excitation during development [45]. Therefore, we next analyzed how a tonic GABAergic conductance (g_GABA_^tonic^) influences g_AMPA_^Thr^ and E_GABA_^Thr^ in a ball model (Fig. 6A), using a g_GABA_^tonic^ between 8.75 fS/cm^2^ and 8.75 nS/cm^2^, corresponding to values from 1/100 to 10000 times of the experimentally determined tonic GABA conductance of 0.875 pS/cm^2^ [52].

**Figure 6.**
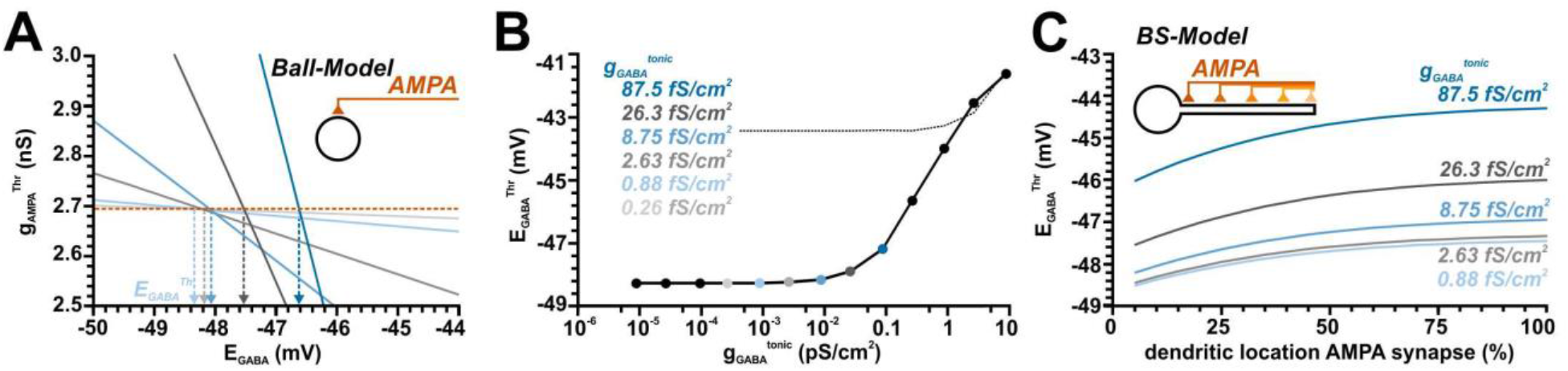
Influence of tonic GABAergic conductances on the AMPA receptor-dependent excitability in a simple ball and a ball-and-stick model. A: Plot of g_GABA_^Thr^ at different E_GABA_. The differently shaded lines represent different tonic g_GABA_ values as indicated in B. The increased slope of the curves with higher g_GABA_^tonic^ illustrates the higher inhibitory effect under this conditions. From the intersection of the plots with the g_GABA_^Thr^ value obtained in the absence of tonic GABA (orange line) the E_GABA_^Thr^ values were determined. B: Plot of E_GABA_^Thr^ determined at different g_GABA_^tonic^. The dashed line represents E_Thr_^ST^. Note that E_GABA_^Thr^ is negative to E_Thr_^ST^ for g_GABA_^tonic^ < ca 3 pS/cm^2^. C: Influence of different dendritic locations of AMPA synapses in a ball-and-stick model on the AMPA receptor-dependent excitability determined for different g_GABA_^tonic^. Note the substantial shift of E_GABA_^Thr^ to positive values with more distant AMPA synapses and the systematic depolarized shift with increasing g_GABA_^tonic^.

These experiments demonstrated, that g_GABA_^tonic^ can attenuate or enhance AP induction by AMPA synapses, depending on E_GABA_. As expected, the slope of the GABAergic influence increased with g_GABA_^tonic^ (Fig. 6A). And as expected, tonic GABAergic conductance enhanced g_AMPA_^Thr^ at hyperpolarized E_GABA_, while smaller g_AMPA_^Thr^ values were required at more depolarized E_GABA_ (Fig. 6A). From the intersection of these g_AMPA_^Thr^ with the basal g_AMPA_^Thr^ (obtained in the absence of tonic GABA), E_GABA_^Thr^ was determined (Fig. 6B). Notably, these E_GABA_^Thr^ were rather constant at ca. −48.3 mV within a wide range of g_GABA_^tonic^, spanning from 0.001 to ca. 10 times the experimentally determined g_GABA_^tonic^ value. Only at very high g_GABA_^tonic^ of > 100 fS/cm^2^ E_GABA_^Thr^ approached E_Thr_^ST^ (which under these conditions was shifted to positive values due to the massively enhanced total membrane conductance). In summary, these results indicate that tonic GABAergic conductances can mediate an excitatory effect even if E_GABA_ was substantially negative to E_AP_^Thr^.

Finally, we investigated how the E_GABA_ of g_GABA_^tonic^ affects the excitation generated by AMPA synapses located along the dendrite in a ball-and-stick model (Fig. 6C). These simulations revealed (i) that E_GABA_^Thr^ was systematically shifted to positive values (closer to E_AP_^Thr^) for distal AMPA synapses and (ii) that E_GABA_^Thr^ was also more positive (and thus closer to E_AP_^Thr^) for larger g_GABA_^tonic^ at all dendritic positions (Fig. 6C). These observations suggest that a tonic GABAergic conductance mediates an excitatory effect even at E_GABA_ that is substantially negative to E_AP_^Thr^, but that an inhibitory effect of tonic GABAergic conductance is higher at distal AMPA-mediated inputs.

## 3. Discussion

Experimental findings indicate that [Cl^−^]_i_ and [HCO^3−^]_i_ are dynamically shifted during early brain development, upon massive GABAergic activity and after pathophysiological insults [10,15,61]. Thus it became evident that GABA can have depolarizing actions [8,13] and this raised the question under which conditions the activation of GABA_A_ receptors can mediate an excitatory effect. Theoretical considerations suggested that GABA_A_ receptor activation permits an inhibitory effect as long as E_GABA_ was below E_Thr_^AP^ [35,36]. However, this consideration just reflects a quasi one-dimensional situation and ignores the temporal and spatial components of GABAergic membrane responses as well as the restriction imposed by the passive membrane properties within more complex neuronal topologies [37–39]. Because the exact role of GABA on the excitation/inhibition threshold is therefore hard to predict from such theoretical assumptions, we performed a detailed in-silico study using a simple neuronal topology and distinct spatiotemporal relations between GABAergic and glutamatergic inputs to evaluate at which E_GABA_ values the net GABA effect switches from inhibitory to excitatory. In these simulations we were able to demonstrate that (i) for GABAergic synapses located close to the AP initiation zone (AIP) the difference between E_GABA_ and E_AP_^Thr^ indeed reliably predicts whether GABA has an excitatory or inhibitory action. (ii) The threshold GABA reversal potential (E_GABA_^Thr^) was in this case close to the E_AP_^Thr^ defined by the maximal subthreshold current injection (E_Thr_^ST^). (iii) E_GABA_^Thr^ was systematically shifted to positive values with increasing distance between the GABA synapse and the AIP. (iv) An excitatory effect of GABA inputs on synchronous AMPA mediated inputs was observed when E_GABA_ was above −44.9 mV, and thus consistently hyperpolarized to E_AP_^Thr^. (v) E_GABA_^Thr^ critically depends on the temporal relation between GABA and AMPA inputs, with a striking excitatory effect on AMPA-mediated inputs appearing after the GABA input. (vi) The spatial relation between GABAergic and AMPA-mediated inputs critically influences E_GABA_^Thr^, with E_GABA_^Thr^ systematically being shifted to values negative to E_AP_^Thr^ for AMPA synapses located proximally to the GABA input. (vii) For tonic GABAergic conductances, E_GABA_^Thr^ was systematically negative to E_AP_^Thr^ over a wide range of g_GABA_^tonic^ values spanning the physiological range. In summary, these results demonstrate that only for very restricted conditions the GABAergic effects switch from excitation to inhibition when E_GABA_ was at E_AP_^Thr^. Under several physiologically relevant conditions, E_GABA_^Thr^ was negative to E_AP_^Thr^, suggesting that GABA can mediate excitatory effects already under these conditions.

It is important to note that in the present study we considered only E_GABA_ as the relevant parameter, which in reality depends not only on [Cl^−^]_i_ but also on [HCO_3_^−^]_i_ [6]. We have chosen this approach to (i) ease the computational load, (ii) because the consideration of two independent variables makes the interpretation of the results more complicated, and (iii) because the relative HCO_3_^−^ conductance of GABA_A_ receptors differs between distinct neuronal subpopulations [6,62,63]. Differences in intracellular fixed charges can also slightly influence the relation between [Cl^−^]_i_, E_Cl_ and the GABAergic driving force [64,65]. In addition, we did not consider that functionally relevant somato-dendritic [Cl^−^]_i_ gradients exists in neurons [11,66] and that GABAergic synaptic activity, alone or correlated to glutamatergic inputs, considerably alters E_GABA_ [49,52,61,67–70]. All of these properties will complicate the prediction of GABAergic response direction, however, for any interpretation of the functional consequences of temporal and spatially dynamic [Cl^−^]_i_ (and [HCO_3_^−^]_i_) gradients, it will be necessary to obtain a major framework to understand how the GABAergic response direction depends on the relation between E_GABA_, E_AP_^Thr^ and spatiotemporal synaptic properties.

The first major result of this in-silico study was the observation that E_GABA_^Thr^ determined for the GABAergic effect on AMPA-mediated inputs was in many cases considerably negative to E_AP_^Thr^, in contrast to the initial theoretical consideration [35,36]. In our experiment we were also able to provide a mechanistic explanation for this observation. First, by using a current-clamp approach we could replicate that the GABAergic depolarization, when isolated from the GABAergic conductance shift, acted excitatory whenever the peak GABAergic depolarization was positive to the RMP, resulting in an E_GABA_^Thr^ of −50.5 mV. This stringent excitatory effect can be easily explained by the fact that in the absence of conductance changes each depolarization brings E_m_ closer to E_AP_^Thr^. Next, we could demonstrate, by providing AMPA-inputs with a defined advance or delay to the GABAergic inputs, a clear bimodal effect of depolarizing GABA responses. In all cases in which the AMPA inputs preceded the GABA input E_GABA_^Thr^ was close to E_AP_^Thr^ (Fig. 4H). Under this condition the AP initiation was under the control of the subsequent GABAergic conductance shift. And under this condition, the GABA_A_ receptor will mediate an inward current, corresponding to a putative excitatory effect, as long as E_GABA_ was positive to E_m_, Thereby, an excitatory effect was induced only if E_GABA_ was above E_AP_^Thr^. However, if the AMPA-mediated inputs occurred after the GABAergic inputs, E_GABA_^Thr^ was systematically shifted to more negative values approximating the RMP of −50.5 mV. This effect can be attributed to the fact that the GABAergic depolarization outlasts the GABAergic conductance shift. Thus, under these conditions the depolarization progressively dominates the effect of GABA, leading to a gradual shift in E_GABA_^Thr^ towards more negative potentials. If the GABAergic conductance can be neglected, each depolarizing shift, i.e. each membrane change depolarized to RMP, contributed to the excitation, leading again to an E_GABA_^Thr^ of −50.5 mV. The impact of the temporal profile of GABAergic conductance change vs. GABAergic depolarization on neuronal excitability has already been experimentally addressed in hypothalamic [39] and neocortical [40] neurons, where the initial phase of a GABA response prevented AP initiation, whereas at later time points of the GABAergic responses AP initiation was facilitated. Despite this clear latency-dependent effect, the reciprocal actions of a depolarization-induced facilitation and a conductance-induced shunting inhibition can also explain why E_GABA_^Thr^ for synaptic inputs was neither at RMP, which would be the case if only the membrane potential shift was relevant, nor at E_Thr_^AP^, which would be the case if E_m_ was only dependent on the actual GABAergic conductance.

In immature neurons, with their slow membrane time constants [71,72], the membrane responses are most probably prone to outlast the membrane conductance for both glutamatergic and GABAergic synaptic inputs. On the other hand, this effect of a prolonged membrane time constant in immature neurons may be partially compensated by the fact, that immature synaptic GABAergic currents show significantly longer decay time constants [72], thereby prolonging the interval in which the shunting inhibitory effects contributes to E_GABA_^Thr^. Another important functional consequence of our results is that the timing between GABAergic and glutamatergic inputs critically determines E_GABA_^Thr^. In this respect classical feedforward as well as recurrent inhibition, with its short latency to excitatory inputs [73], will impose a rather strict inhibition even at depolarizing GABAergic conditions as long as E_GABA_ is maintained below E_Thr_^AP^. Thus this kind of inhibitory control would be rather stable upon activity dependent shifts in E_GABA_ [49,61,67,68,74]. On the other hand, for GABAergic inputs that are not temporally correlated with the excitatory inputs, e.g. during blanket inhibition, it must be considered that E_GABA_^Thr^ can be negative to E_AP_^Thr^, and thus may mediate a less stable inhibition that is more sensitive to ionic plasticity.

The second major result of this in-silico study was the observation, that the spatial relation between GABAergic and AMPA inputs also critically affects E_GABA_^Thr^. As expected, our simulation revealed that the inhibitory effect, as quantified by Δg_AMPA_^Thr^, of proximal GABAergic synapses are stronger than that of distally located ones. The Δg_AMPA_^Thr^ values were substantially larger for AMPA synapses located distally to the GABA synapse, indicating that a GABA input can shunt EPSPs from distally located excitatory synapses, as suggested from in-vitro and in-silico experiments [40]. For proximally located GABA synapses we could observe that E_GABA_ showed only little dependency on the location of the AMPA-mediated inputs. In these cases, E_GABA_^Thr^ amounted to ca. −46 mV, suggesting that both, shunting and depolarizing effects contribute to the impact of GABA on the excitability. In contrast, we observed for distally located GABA synapses a strong dependency of the location of AMPA-mediated inputs on E_GABA_^Thr^. For such distal GABA synapse locations a negative E_GABA_^Thr^ close to −50 mV was observed at proximal AMPA synapses, which reflects the fact that with this configuration only the electrotonically propagating GABAergic depolarization has an effective influence on the AMPA-mediated depolarization, while the GABAergic conductance shift acts more locally. For co-localized GABA and AMPA synapses at the distal end of the dendrite E_GABA_^Thr^ approximated E_AP_^Thr^ at ca. −43 mV, indicating that here the effect of GABA was mediated mainly by membrane shunting. Intriguingly the “slope” of E_GABA_^Thr^ was steeper for AMPA synapses in the dendritic segment proximal to the GABA synapse. The slope became shallower for the segment distal from the GABA synapse. This observation indicates that for all AMPA synapses distal to the GABA synapse a substantial fraction of the synaptic currents were shunted by the GABAergic conductance before they can affect AP initiation in the soma. In contrast, for all AMPA synapses located proximal to the GABA synapse the shunting effect was diminished with increasing distance between both synapses, whereas the electrotonically propagating depolarization maintained a more stable excitatory influence and thereby shifted E_GABA_^Thr^ towards the RMP. Thus the results of our experiments suggest an additional mechanism that contribute the putative excitatory GABAergic effect of dendritic GABA inputs [40], in addition to the existence of stable or dynamic somato-dendritic [Cl^−^]_i_ gradients [75,76].

These in-silico observations indicate that perisomatic inhibition, which is the dominant form for the classical feedback and feedforward inhibition mediated by parvalbumin-positive interneurons [77,78], can maintain a stable inhibitory effect regardless of the site of glutamatergic inputs and ionic plasticity. On the other hand, the impact of GABAergic synapses located in the dendritic periphery, e.g. by the hippocampal O-LM interneurons [79] or neocortical Martinotti interneurons [80], will critically depend on the location of the excitatory glutamatergic inputs and can putatively mediate an excitatory impact on AMPA synapses close to the soma at slightly depolarizing E_GABA_.

In addition, our results indicate that for small to moderate tonic GABAergic conductance E_GABA_^Thr^ was systematically more negative than E_AP_^Thr^, which suggests that even at rather moderate depolarizations tonic GABAergic currents can mediate an excitatory effect. Only at higher g_GABA_^tonic^ the E_GABA_^Thr^ approaches E_AP_^Thr^. The results of this simulation replicate the findings of a previous in-vitro study, that demonstrated excitatory effects of depolarizing tonic GABAergic responses at low conductances, whereas at higher conductances a stable inhibition was imposed [81]. Our results are also in line with the excitatory effects of extrasynaptic GABA_A_ receptors in the immature hippocampus [45]. In our simulations E_GABA_^Thr^ remained stable at about −48.3 mV for g_GABA_^tonic^ smaller than ca. 10^−2^ pS/cm^2^, which is close to the passive membrane conductance g_pas_ of 0.0128 pS/cm^2^. We assume that below this value the shunting effects caused by g_GABA_^tonic^ were negligible to the background conductance g_pas_ and thus did not considerably contribute to the shunting of EPSCs. Only if g_GABA_^tonic^ exceeded g_pas_ a relevant additional inhibitory component was imposed by the GABAergic conductances and thus E_GABA_^Thr^ converged towards E_AP_^Thr^.

Another conclusion that could be drawn from our study is that some attention should be taken to the method used to detect the AP threshold. Obviously there is, despite the frequent use of this descriptive parameter, no consensus on the definition of AP threshold [42]. Therefore, we used in this in-silico study four different, established methods for E_AP_^Thr^ detection. Our in-silico experiments demonstrated that the AP threshold value determined from a fixed threshold of dV/dt [44,50], from the first positive peak in d^3^V/dt^3^ [51], and from linear regression of the AP upstroke [43] were comparable at potentials of ca. −34 mV to −37 mV. In contrast, substantially negative values of −43 mV were determined if E_AP_^Thr^ was defined as the maximal potential that did not result in AP triggering (E_Thr_^ST^). The difference in the results of these methods can be easily explained by the fact that E_Thr_^ST^ represents a quasi-stationary value (dV/dt close to 0) that is just insufficient to trigger the entry to the Hodgkin cycle. On the other hand, the first three E_AP_^Thr^ values represent distinct states during the dynamic events in the initial AP phase. The fact that in our simulations E_GABA_^Thr^ for only GABAergic inputs indeed approximated E_Thr_^ST^ can be related to the fact that the excitation threshold for GABAergic inputs was also determined under quasi-stationary conditions. For the influence of GABA on synaptic AMPA-mediated inputs the excitation threshold was determined in the interval between the onset of the GABA inputs and the duration at which 63% of the peak depolarization was obtained. Thus, for the relevant traces that distinguished between subthreshold and suprathreshold AMPA inputs, dV/dt was considerable small and thus the AP threshold was also determined under quasi stationary conditions. Under physiological conditions random fluctuation in E_m_ will most probably limit the difference between E_Thr_^dVdt^, E_Thr_^d3^, E_Thr_^IS^, and E_Thr_^ST^. In any way, while addition of membrane noise to the in-silico models and/or a different methodological definition of the excitation threshold for GABA- and AMPA-mediated inputs would probably change the absolute values for E_GABA_^Thr^ and E_AP_^Thr^, it would not substantially interfere with the main observation of this study, that E_GABA_^Thr^ is for many physiologically relevant situations negative to E_AP_^Thr^.

In conclusion, this simulation indicates that, in addition to the influence of short-term and long-term ionic plasticity, the uneven distribution of [Cl^−^]i gradients within individual cells and the effects of tonic and phasic inhibition [10,11,61,67], the observed spatial and temporal constraints on the E_GABA_ to E_AP_^Thr^ relation imposes another level of complexity to the dynamic properties of GABAergic inhibition/excitation. While on one hand our results support the textbook knowledge that GABA mediates a stable inhibition as long as hyperpolarizing membrane responses are evoked (or [Cl^−^]_i_ is sufficiently low), on the other hand the altered [Cl^−^]_i_ homeostasis in early development and several neurological conditions like trauma, stroke or epilepsy [11,12,30,31], can impact the level of inhibitory control already upon moderate [Cl^−^]_i_ changes in a complex way.

## 4. Materials and Methods

### 4.1. Electrophysiological procedures

#### 4.1.1. Slice preparation

All experiments were conducted in accordance with EU directive 86/609/EEC for the use of animals in research and the NIH Guide for the Care and Use of Laboratory Animals, and were approved by the local ethical committee (Landesuntersuchungsanstalt RLP, Koblenz, Germany). We made all efforts to minimize the number of animals and their suffering. Newborn pups of postnatal days [P] 4-7 were obtained from time pregnant C57Bl/6 mice (Janvier Labs, Saint Berthevin, France) housed in the local animal facility at 12/12 day/night cycle and ad libitum access to food and water. The mouse pups were decapitated in deep enflurane (Ethrane, Abbot Laboratories, Wiesbaden, Germany) anaesthesia, their brains were quickly removed and immersed for 2-3 min in ice-cold standard artificial cerebrospinal fluid (ACSF, 125 mM NaCl, 25 mM NaHCO_3_, 1.25 mM NaH_2_PO_5_, 1 mM MgCl_2_, 2 mM CaCl_2_, 2.5 mM KCl, 10 mM glucose, equilibrated with 95% O_2_ / 5 % CO_2_, osmolarity 306 mOsm). Four hundred μm thick horizontal slices including the hippocampus were cut on a vibratome (Microm HM 650 V, Thermo Fischer Scientific, Schwerte, Germany) and subsequently stored in an incubation chamber filled with oxygenated ACSF at room temperature for at least 1h before they were transferred to the recording chamber.

#### 4.1.2. Patch-clamp recordings

Whole-cell patch-clamp recordings were performed at 31 ± 1 °C in a submerged-type recording chamber attached to the fixed stage of a microscope (BX51 WI, Olympus). Pyramidal neurons in the stratum pyramidale of the CA3 region were identified by their location and morphological appearance in infrared differential interference contrast image. Patch-pipettes (5-12 MΩ) were pulled from borosilicate glass capillaries (2.0 mm outside, 1.16 mm inside diameter, Science Products, Hofheim, Germany) on a vertical puller (PP-830, Narishige) and filled with the pipette solutions (86 mM K-gluconate, 44 mM KCl, 4 mM NaCl, 1 mM CaCl_2_, 11 mM EGTA, 10 mM K-HEPES, 2 mM Mg2-ATP, 0.5 mM Na-GTP, pH adjusted to 7.4 with KOH and osmolarity to 306 mOsm with sucrose). In few experiments 40 mM KCl were replaced with 40 mM K-gluconate. Signals were recorded with a discontinuous voltage-clamp/current-clamp amplifier (SEC05L, NPI, Tamm, Germany), low-pass filtered at 3 kHz and stored and analyzed using an ITC-1600 AD/DA board (HEKA) and TIDA software. All voltages were corrected post-hoc for liquid junction potentials of −8 mV for a [Cl^−^] of 10 mM and −5 mV for 50 mM [20]. Input resistance and capacitance were determined from a series of hyperpolarizing current steps. Action potentials (AP) were induced by a series of depolarizing current steps. For averaging of AP wave forms the first AP from traces that showed a series of APs were used.

### 4.2. Compartmental modeling

The compartmental modeling was performed using the NEURON environment (neuron.yale.edu). The source code of models and stimulation files used in the present paper can be found in ModelDB (http://modeldb.yale.edu/267062; reviewer password is “GABA”). For compartmental modelling we used either a simple ball (soma diameter = 43 μm) or a ball and stick model (soma with d=43 μm, linear dendrite with l=1000 μm, diameter 1 μm, and 301 nodes). In both models a passive conductance (g_pas_) with a density of 1.28*10^−5^ nS/cm^2^ and a reversal potential (E_pas_) of −50.5 mV was distributed, which resulted for the ball-and-stick model in passive membrane properties that were comparable to the properties of recorded pyramidal CA3 neurons.

Because it was not possible to generate a reasonable sharp AP onset with a standard Hodgkin-Huxley (HH) model and since we are particularly interested in the AP threshold properties, we heuristically developed a modified Markov model, massively simplified from published Markov models [82,83] to simulate the AP with a considerably precision. For this modified Markov model we consider only 3 different states for the Na+ channels [84]:

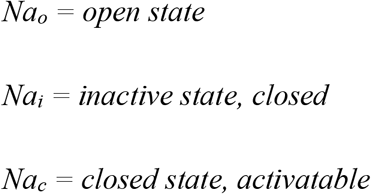

We restricted transitions between these states to Na_c_→Na_o_, Na_o_→Na_i_, Na_i_→Na_c_, Na_c_→Na_i_.

The transition rate Na_c_→Na_o_ is only voltage dependent as described by a Bolzmann function:

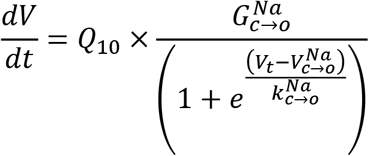

The transition Na_o_→Na_i_ is simulated by a voltage dependent kinetic rate described by a Bolzmann equation with is operational after a defined delay period plus a constant voltage-independent term

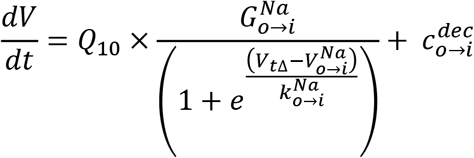

The transition rates Na_c_→Na_i_ and Na_i_→Na_c_ are described again by simple Bolzmann functions as follows:

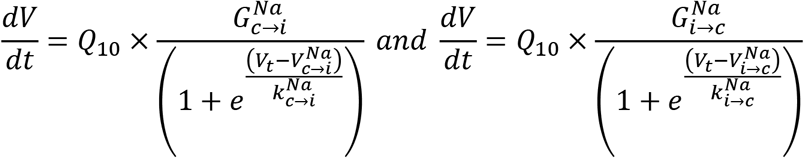

In addition, we implemented a simple two state modified Markov model for the delayed rectifier K^+^ current, with the K_c_→K_o_ transition rate described by a Bolzmann equation with an operational delay period as follows:

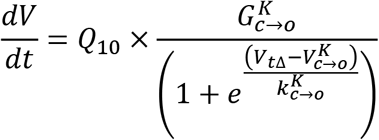

And the K_o_→K_c_ transition rate described by a Bolzmann equation with an operational delay period:

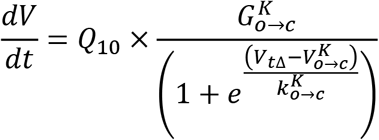

All states for the Na^+^ and K^+^ channels are normalized at each iteration step as follows:

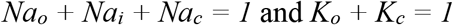

The Na^+^ current was given according to Ohms law as:

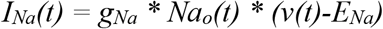

And the K^+^ current was as:

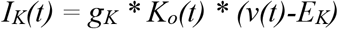

All parameters were optimized by stepwise approximation to obtain a sufficient fit to the average experimentally determined AP trace, which was quantified by minimizing the root of the summarized squared errors according to the following error weight function:

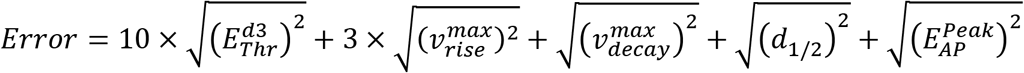

This error function was used with the rationale to put special emphasise for the fitting routine to the dynamic properties at E_AP_^Thr^. The used parameters are given in Table 2.

**Table 1.** Parameters used for the sigmoidal fit of the E_GABA_ to g_GABA_^Thr^ relationship and the resulting E_GABA_^Thr^.

| **Phasic GABA - distributed GABA inputs** |  |  |  |  |  |
| --- | --- | --- | --- | --- | --- |
| **g<sub>GABA</sub>** | **p<sub>Max</sub>** | **E<sub>0</sub>** | **s** | **k** | **E<sub>GABA</sub><sup>Thr</sup>** |
| 0.0158 pS | 1 pS | -36.0 mV | -20.4 mV | 0.75 | -43.05 mV |
| 0.0789 pS | 1 pS | -37.7 mV | -2.5 mV | 0.28 | -43.35 mV |
| 0.3945 pS | 1 pS | -41.6 mV | -0.47 mV | 0.21 | -43.06 mV |
| **Phasic GABA - proximal GABA inputs** |  |  |  |  |  |
| **g<sub>GABA</sub>** | **p<sub>Max</sub>** | **E<sub>0</sub>** | **s** | **k** | **E<sub>GABA</sub><sup>Thr</sup>** |
| 0.0158 pS | 1 pS | -38.5 mV | -19.3 mV | 0.80 | -43.32 mV |
| 0.0789 pS | 1 pS | -38.2 mV | -2.5 mV | 0.30 | -43.41 mV |
| 0.3945 pS | 1 pS | -41.8 mV | -0.49 mV | 0.24 | -43.12 mV |
| **Phasic GABA - distal GABA inputs** |  |  |  |  |  |
| **g<sub>GABA</sub>** | **p<sub>Max</sub>** | **E<sub>0</sub>** | **s** | **k** | **E<sub>GABA</sub><sup>Thr</sup>** |
| 0.0158 pS | 1 pS | -39.5 mV | -22.3 mV | 0.85 | -43.10 mV |
| 0.0789 pS | 1 pS | -37.5 mV | -2.8 mV | 0.30 | -43.34 mV |
| 0.3945 pS | 1 pS | -41.45 mV | -0.6 mV | 0.24 | -43.09 mV |
| **Tonic GABA** |  |  |  |  |  |
| **g<sub>GABA</sub>** | **p<sub>Max</sub>** | **E<sub>0</sub>** | **s** | **k** | **E<sub>GABA</sub><sup>Thr</sup>** |
| 0.0875 pS | 1.0 pS | -52.70 mV | -350.00 mV | 1 | -35.90 mV |
| 0.1750 pS | 1.0 pS | -49.60 mV | -226.00 mV | 1 | -38.75 mV |
| 0.2625 pS | 1.0 pS | -47.60 mV | -148.00 mV | 1 | -40.50 mV |
| 0.4375 pS | 1.0 pS | -46.10 mV | -87.00 mV | 1 | -41.93 mV |
| 0.8750 pS | 1.0 pS | -44.90 mV | -39.50 mV | 1 | -43.00 mV |
| 1.7500 pS | 1.0 pS | -24.30 mV | -17.50 mV | 0.5 | -43.41 mV |
| 2.6250 pS | 1.0 pS | -32.55 mV | -10.50 mV | 0.5 | -43.42 mV |
| 4.3750 pS | 1.0 pS | -36.95 mV | -6.20 mV | 0.45 | -43.59 mV |
| 8.7500 pS | 1.0 pS | -39.57 mV | -2.55 mV | 0.38 | -43.58 mV |
| 17.5000 pS | 1.0 pS | -41.34 mV | -1.17 mV | 0.33 | -43.55 mV |
| 26.2500 pS | 1.0 pS | -41.78 mV | -0.74 mV | 0.29 | -43.41 mV |
| 43.7500 pS | 1.0 pS | -42.26 mV | -0.48 mV | 0.32 | -43.20 mV |
| 87.5000 pS | 1.0 pS | -42.35 mV | -0.22 mV | 0.3 | -42.82 mV |

**Table 2.**
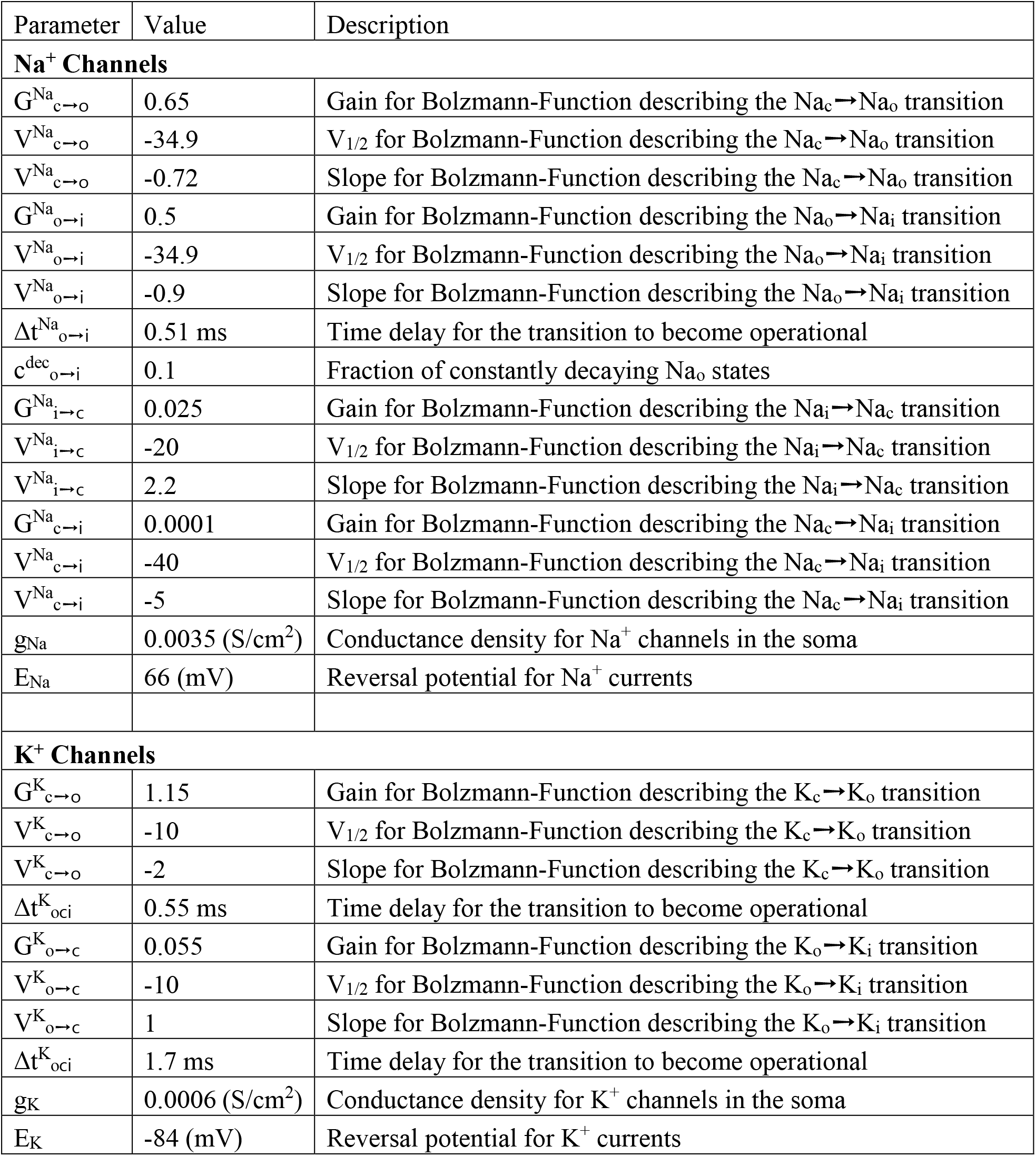
Parameters used for the modified Markov model.

AMPA synapses were modeled by an Exp2Syn point process using a reversal potential of −12 mV, a tau1 value of 0.1 ms and a tau2 value of 11 ms, in accordance with the experimentally determined value [49]. GABA synapses were modeled by an Exp2Syn point process using a tau1 value of 0.1 ms and a tau2 value of 37 ms, in accordance with the experimentally determined value [49]. The reversal potential of the GABAergic synapses was the main variable of interest in these simulations. For tonic GABAergic currents a constant membrane conductance was distributed over all membrane with conductance densities and reversal potentials as given in the results part.

For the determination of g_GABA_^Thr^ we used an iterative approach where g_GABA_ was first increased by 1 nS steps until an AP was induced within an interval of 800 ms after the GABA input. Subsequently g_GABA_ was decreased by 0.33 nS steps until the AP vanished, followed again by an increase in g_GAB_A by 0.1 pS until the AP reappeared. This alternating sequence was repeated 6 times using a g_GABA_ of 1/10 for each subsequent round. In these experiments E_AP_^Thr^ was defined as the peak voltage of the last subthreshold sweep.

A similar approach was also used to determine g_AMPA_^Thr^. Here g_AMPA_ was initially increased by 0.01 nS steps until an AP was induced. The analysis interval was in all sweeps set to the interval between stimulus onset and the time point when the AMPA-mediated depolarization, determined in the absence of an AP mechanism, decreased to 63% of the peak amplitude. Subsequent *g_AMPA_* was decreased by 3.3 pS until the AP disappears, followed by 6 rounds of alternating increasing/decreasing *g_AMPA_* steps, with *g_AMPA_* step values decreasing by 1/10 for each round.

## Author Contributions

Conceptualization: WK; Electrophysiological investigation: AL, Formal analysis, AL and WK; Modeling: WK: Writing WK, AL, and HJL.

## Suppl. Data

**Suppl. Fig. 1.**
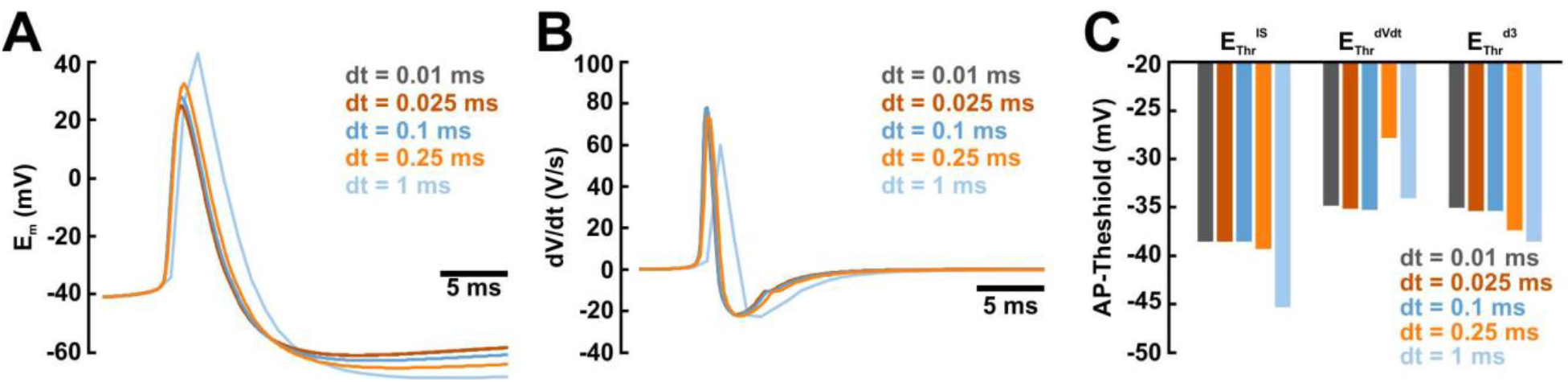
Characterization of AP properties using different dt values for the simulation. A: Simulated voltage traces using different dt as indicated in the plot. Note the divergence of AP shape at larger dt values. B: Rate of E_m_ changes during an action potential. C: Typical E_AP_^Thr^ values determined with 3 different algorithms on the traces obtained at different dt. Note that all E_Thr_^IS^, E_Thr_^dV/dt^ and E_Thr_^d3^ remained stable for a dt ≤ 0.1 ms.

